# Antenatal maternal anxiety modulates the BOLD response in 20-year old adolescents during an endogenous cognitive control task

**DOI:** 10.1101/087817

**Authors:** Maarten Mennes, Bea R.H. Van den Bergh, Stefan Sunaert, Lieven Lagae, Peter Stiers

## Abstract

Evidence is building for an association between the level of anxiety experienced by a mother during pregnancy and the cognitive development of her offspring. The current study uses fMRI to examine whether there is an association between prenatal exposure to maternal anxiety and brain activity in 20 year old adolescents. In line with previous results of this follow-up study, it was found that adolescents of mothers reporting high levels of anxiety during weeks 12–22 of their pregnancy had a different pattern of decision making in a Gambling paradigm requiring endogenous cognitive control compared to adolescents of mothers reporting low to average levels of anxiety during pregnancy. Moreover, the blood oxygenation level dependent (BOLD) response in a number of prefrontal cortical areas was modulated by the level of antenatal maternal anxiety. In particular a number of right lateralized clusters including inferior frontal junction, that were modulated in the adolescents of mothers reporting low to average levels of anxiety during pregnancy by a task manipulation of cognitive control, were not modulated by this manipulation in the adolescents of mothers reporting high levels of anxiety during pregnancy. These results provide a neurobiological underpinning for our previous hypothesis of an association between a deficit in endogenous cognitive control in adolescence and exposure to maternal anxiety in the prenatal life period.

## 1 Introduction

Previous phases of our longitudinal study into the association between maternal anxiety during pregnancy and the development of the offspring have provided substantial evidence showing that high levels of anxiety experienced by a mother during pregnancy influence the cognitive development of her offspring up into adolescence. In 15 and 17 year old adolescents specific behavioral neurocognitive measures were used, linking antenatal maternal anxiety to a deficit in endogenous cognitive control (Van den Bergh et al., 2005, 2006; Mennes, Stiers, Lagae, & Van den Bergh, 2006). This function refers to the ability to generate control over decisions, strategies, conflicting information and so on from within oneself without external signals triggering this control. By coupling these findings to evidence of brain imaging studies that used similar measures we hypothesized the functioning of the orbitofrontal cortex to be most affected by antenatal maternal anxiety (Mennes et al., 2006). However, this hypothesis was only indirectly raised as no information on the actual brain functioning of the investigated sample was available.

A first attempt to gain more insight in the actual brain functioning of the adolescents of our study sample was made in the follow-up phase at age 17 by measuring the electrical brain activity of the adolescents using EEG. The hypothesis of an impairment in endogenous cognitive control was further delineated by contrasting a Go/Nogo task assessing exogenous cognitive control with a Gambling paradigm assessing endogenous cognitive control. Confirming our hypotheses, there was no evidence for a relationship between antenatal maternal anxiety and the behavioral and ERP results of the Go/Nogo task. In the Gambling paradigm on the other hand, the level of antenatal maternal anxiety modulated an early frontal peak (P2a) (Mennes, Van den Bergh, Lagae, & Stiers, 2009). These results provided the first evidence for an association between antenatal exposure to maternal anxiety and actual brain functioning.

Although ERPs are a valid measure of brain activity they do not allow a precise localization of the activity in terms of specific areas of the brain that contribute to the activity measured at the scalp. Functional Magnetic Resonance Imaging (fMRI) on the other hand, is capable of providing information about the functionality of the brain with a good spatial resolution. Here we used fMRI to investigate the association between antenatal maternal anxiety and endogenous cognitive control as measured with our Gambling paradigm.

This study will explore whether the hemodynamic BOLD response measured in 20 year old adolescents can be related to the level of anxiety experienced by their mother during weeks 12–22 of their pregnancy. Based on the previous results in the same sample we hypothesize such di erences to be situated in the prefrontal cortex, and more specifically in orbitofrontal cortex (Van den Bergh et al., 2005, 2006; Mennes et al., 2006). We will investigate the adolescents with the Gambling paradigm used before with ERP in the same research sample (Mennes et al., 2009). In this paradigm participants have to assess risks, make decisions, monitor their total scores and deal with gains and losses, all requiring endogenous cognitive control. As prefrontal cortex and its connected circuitry is regarded the control center for such complex executive behavior we expect different prefrontal areas to be modulated by this manipulation (Miller & Cohen, 2001). In addition, recent imaging studies using task-switching paradigms identified a region labelled inferior frontal junction, at the junction between inferior frontal sulcus and inferior precentral sulcus, as a key region in endogenous cognitive control, next to mid-dorsolateral prefrontal cortex and medial-frontal regions (Brass & von Cramon, 2004; Brass, Derrfuss, Forstmann, & von Cramon, 2005; Forstmann, Brass, Koch, & von Cramon, 2005). These regions might also be modulated by an additional manipulation implemented in the Gambling paradigm. By manipulating the individual trial characteristics we made the required decisions either more exogenous (trial-based) or more endogenous (subject-based). The behavioral results gathered with the Gambling paradigm at age 17 yielded an association between this manipulation and the level of anxiety experienced by their mothers during pregnancy. The ERPs on the other hand, showed no relationship between antenatal maternal anxiety and the exogenous/endogenous control manipulation, although they yielded a main effect of antenatal maternal anxiety on the frontal P2a peak. Due to the discrepancies in temporal and spatial resolution between ERPs and fMRI it is plausible that we will observe modulations in the BOLD response related to both antenatal maternal anxiety and the exogenous/endogenous manipulation.

## 2 Methods

### 2.1 Participants

Previous results of the longitudinal research project this study is part of yielded more consistent results in boys compared to girls. This might be due to the fact that the assessed cognitive functions are more likely to be associated with antenatal maternal anxiety in males, compared to a higher chance for mood-related disorders in girls (Van den Bergh, Van Calster, Smits, Van Huffel, & Lagae, 2008). For this reason and the fact that the high anxiety group comprised only five girls, we included only boys in the current study.

Eighteen adolescent boys were selected based on the anxiety grouping at age 17. At that age the adolescents were divided in a low-average and high anxiety group based on the anxiety scores of their mothers during weeks 12–22 of their pregnancy (see Mennes et al., 2006). Ten boys ware part of the high anxiety group and except for one boy who could not be contacted and one boy who wore braces, all were included in the high anxiety group of the current study (*n* = 8). The data of the adolescents in the high anxiety group were compared to those of 10 adolescents of the low-average anxiety group. The adolescent boys of the low-average anxiety group were selected to match the cognitive abilities of the adolescents in the high anxiety group.

All participants were 20 years old and were born in the same hospital between 36 and 41 weeks of gestation with a mean birth weight of 3378 grams (SD = 639) and 5 minute Apgar scores of 9 or 10. The local ethical committee for experiments on human subjects approved the study. All participants were clearly informed about the scanning procedures and gave their written informed consent.

Mean PIQ in the low-average group was 99.67 (SD = 9.95), compared to 98.25 (SD = 8.65) for the adolescents in the high anxiety group. This difference was not significant, proving that both groups were appropriately matched (*t*_15_ = -.31, *p* = .76).

One boy of the low-average anxiety group was excluded because his performance on the gambling task was well outside the range of the other boys in the low-average anxiety group. Therefore the final low-average anxiety group included nine adolescent boys (*n* = 9), compared to eight boys in the high anxiety group (*n* = 8).

### 2.2 Gambling Paradigm

A complete description of the Gambling paradigm can be found in Mennes et al., 2008 (Mennes, Wouters, Van den Bergh, Lagae, & Stiers, 2008). The same paradigm was used in the current study. Figure 1 shows a stimulus sequence. A two-sided colored bar appeared on the screen. Participants could gamble on the side they thought a token was hidden underneath by pressing a button according to the chosen side. A correct gamble resulted in winning a number of points that was shown above the stimulus bar (ranging from 10 to 100). An incorrect gamble resulted in losing a number of points shown below the bar (ranging from 0 to 100). However, subjects were also given the opportunity to refrain from making a gamble. If participants felt insecure about their decision, or if they found the gain too low or the loss too high, they could choose not to gamble. In that case they just had to wait for the stimulus to disappear; no key press had to be given. Refraining from gambling always resulted in earning 20 points. All participants received a 100 point credit to start with and were verbally motivated to earn as much points as possible.

**Figure 1:**
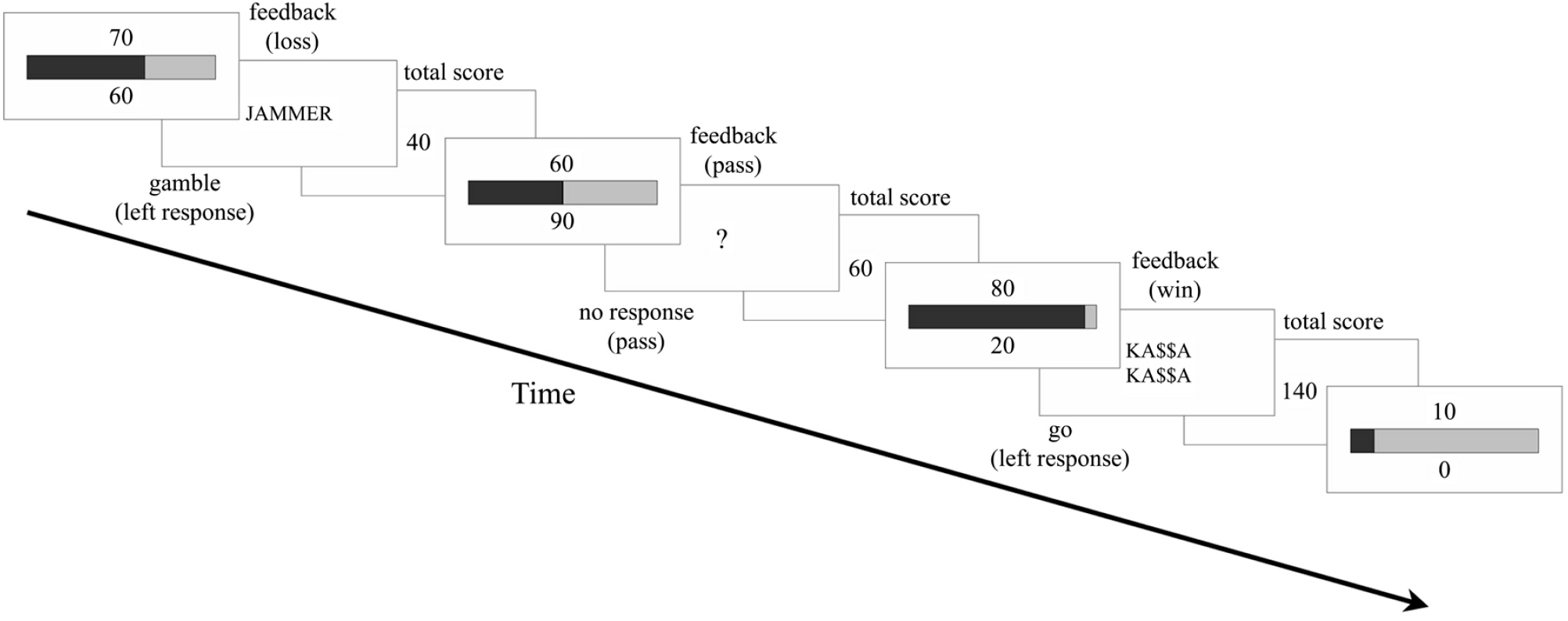
Stimulus sequence of the Gambling paradigm. On each trial participants were shown a proportionally divided colored bar (blue and yellow). Participants could gamble on the location of a hidden token. A correct gamble was rewarded the points shown above the bar, an incorrect gamble resulted in a loss of the points shown below the bar. Participants could also choose not to gamble and settle for a small reward (20 points), as shown in trial 2 of this sequence. After each stimulus bar feedback was given and the total score was updated and shown. (Jammer: Too bad; Ka$$a Ka$$a: ‘ka-ching’).

Two changes were made to the paradigm to optimize it for use in fMRI. First, in 33% of the trials the feedback and total score stimuli were replaced by dummy trials consisting of the gray background screen with a fixation cross in the middle. This was done to avoid a coupling of the BOLD response between the gambling and the outcome stimuli. The dummy trials were randomly mixed with normal feedback trials. Second, the duration of the decision and feedback phase of the trials was randomly jittered in steps of 100 msec. The decision phase lasted between 3800 and 5800 msec. The gambling stimulus itself remained visible for 3500 msec. The length of the feedback phase varied between 2200 and 4800 msec. The feedback stimulus was visible for 900 msec, after an interval of 100 msec the totalscore was shown for 900 msec.

Four decision trial conditions could be dissociated in this paradigm. While the whole paradigm requires endogenous cognitive control, we dissociated two trial types that were more exogenous in nature, and two that were more endogenous. First, there were clearly favorable trials that always led participants to a gamble. This trial type was defined by a win proportion ⩾ 80% and a gain over 20 points. These trials were labeled GO trials. Second, there were trials that should always lead to a pass. This trial condition comprised trials with a gain of only 10 or 20 points, regardless of the proportional division of the colored bar. In these trials gambling was always disadvantageous as participants were certain of a 20 point gain when passing. Therefore these trials were labeled NOGO trials. Both the GO and NOGO trials were considered to be exogenous trials as it is the stimulus that signals the participant on the most appropriate action. The third and fourth trial condition included trials with a proportional division between 50% and 75%, and a gain exceeding 20 points. These characteristics are not clearly pro gambling or pro inhibition. Instead it is not immediately clear to the participants which response should be given. The final decision in these trials will be guided by the interpretation of different kinds of information such as previous experiences, the gain or gain/loss ratio, the total score, or the moment during the task (e.g., at the beginning or near the end). Based on the chosen response, both a GAMBLE and PASS trial condition were defined. These trials are considered endogenous because the participants themselves decide on the most appropriate action without any cue from the stimulus. As such an identical trial sometimes leads to a gamble but other times to a pass depending on the decision of the participant.

All participants completed a practice run of 20 trials before entering the MR environment. During the fMRI session all participants completed four runs of the task. Each run contained 50 gamble trials, resulting in a total of 200 trials for each participant.

### 2.3 fMRI Data Acquisition

Data were acquired on a 3.0-T MR system (Achieva, Philips, Best, the Netherlands) with an eight-channel phased-array head coil. Functional images were acquired using a T2*-weighted gradient echo (GE) echo planar imaging (EPI) sequence (TR = 1950 ms, TE = 33 ms; flip angle = 90°; field of view = 240 × 240 mm^2^; acquisition matrix = 80 × 78; acquired voxel size = 3 × 3.05 × 4.5 mm^3^; 28 4,5 mm axial slices; EPI factor = 61; SENSE reduction factor = 1.4). For each run of the Gambling task 215 dynamic scans were acquired (total acquisition time = 7 min 13 sec). For anatomical reference one high-resolution 3D-TFE T1-weighted structural scan was acquired for each subject (acquisition matrix = 256 × 182; field of view = 250 × 180 mm^2^; flip angle = 8° ; TR = 9.725 ms; TE = 4.6 ms; acquired voxel size = 0.98 × 0.98 × 1 mm^3^; 230 1 mm coronal slices; SENSE reduction factor = 1.4). Stimuli were presented using the Eloquence fMRI system (InvivoMDE, MRI Devices Corporation Inc., Orlando, FL, USA).

### 2.4 Statistics

#### Confounding variables

To establish whether maternal anxiety measured during weeks 23–31 and 32–40 of pregnancy and maternal anxiety measured at each of the postnatal follow-up phases of our study should be included as covariates in the analyses, a prior correlation analysis was performed. This revealed no significant correlations between the dependent variables (gambling contrast value, totalscore) and maternal anxiety during the other pregnancy periods or postnatal maternal anxiety, nor between these confounding variables and maternal anxiety measured during weeks 12–22 of pregnancy, which is the main experimental variable. Therefore, these covariates were not included in the analyses. The adolescents’ intelligence score was also not included as the groups had been matched on intelligence test scores prior to inclusion (as indicated above).

#### Behavioral Data Analysis

Due to its longitudinal design, the number of subjects included in the current phase of the study is rather limited. As there was no homogeneity in the variances of the dependent variables (assessed with Levene’s Test), even after log-transforming these variables to improve normal distribution, we used randomization statistics. We decided to use this procedure instead of traditional parametric statistical tools to assess the effect of anxiety on the untransformed behavioral measures of the Gambling task because these methods are not bound to any underlying parametric assumption. The data for each subject in each of the two anxiety groups were entered in two vectors and the difference between the two means was calculated. After pooling both vectors, two new samples each the size of the previous vectors, were randomly drawn from the total sample and the difference between the means of these two new vectors was calculated. This was repeated 100.000 times, resulting in 100.000 differences. The difference between the original vectors was regarded significant when the probability of observing that difference was less than 5% when assessing the distribution of the differences calculated between the randomly drawn samples.

#### fMRI Analyses

Data were analyzed off-line using SPM5 software (Wellcome Department of Imaging Neuroscience, London, UK). All functional images were slice-time corrected and spatially realigned to the first volume of the first run. After coregistering the functional images to the anatomical image, they were spatially normalized to a custom-made T1 brain in the standard space of the Montreal Neurological Institute. The normalized functional images were resliced into 2 mm^3^ isotropic voxels and were spatially smoothed with a 6 mm full-width at half-maximum Gaussian kernel.

*First*, we wanted to identify prefrontal areas that are related to decision making upon presentation of the gambling stimulus. Identifying such task or condition specific regions will allow interpreting antenatal maternal anxiety related BOLD modulations in relation to the underlying decision-related cognitive processes. Prior to a second-level analysis assessing the effects of the included conditions, a first level analysis was performed on the data of each the nine adolescent boys of the low-average group. The GO, NOGO, GAMBLE and PASS trial conditions were modeled with a canonical haemodynamic response function (HRF) including time and dispersion derivatives. Next to the four decision trial conditions, negative, positive and neutral feedback were also modeled when present (i.e. some participants managed to finish runs without losing and thus also without receiving negative feedback). The six movement parameters of each session were included as covariates in the analysis. Based on this analysis the percentage of signal change was calculated for each of the four decision conditions (GO, NOGO, GAMBLE, and PASS). These maps of the percentage of signal chance in each condition were then used in a second level 2-by-2 random-effects factorial analysis including the factors go/nogo and exogenous/endogenous. These factors were not independent because they comprised repeated measures from the same participants. The 2-by-2 design effectively consisted of four cells each including one of the four decision trial conditions of the Gambling task (go/exogenous: GO; nogo/exogenous: NOGO; go/endogenous: GAMBLE; nogo/exogenous: PASS). Significance for this analysis was assessed at false discovery rate (FDR) level with *p* < .05. Clusters were only assessed if they exceeded 50 voxels in size and were located anterior of central sulcus.

In a *second* analysis we assessed the influence of antenatal maternal anxiety on the BOLD response of the adolescents using a fixed-effects analysis including all four runs of each of the 17 participants. The fixed-effects model, providing more degrees of freedom and statistical power, was chosen due to the limited number of subjects in both groups. As for the individual first level analyses described above, all four decision conditions and the three feedback conditions were modeled with a canonical HRF including time and dispersion derivatives. Again the six movement parameters were included as covariates of no interest for each run. An *F*-contrast for comparing the low-average and high anxiety groups was calculated as a disjunction-contrast including effects of either condition. This means that the conditions were not pooled together but the calculated contrast included differences between the low-average and high anxiety groups in the GO *or* NOGO *or* GAMBLE *or* PASS condition. Significance threshold was set at an alpha level of .05 corrected for multiple comparisons (FWE). Only clusters including more that 50 significant voxels were interpreted. Next, significant clusters from this analysis were used as regions of interest (ROI) in separate 2-by-2 factorial ROI based group analyses. For each ROI the percentage of signal change was calculated for each participant and compared in a repeated measures ANOVA, where participants were treated as a random factor. The model included group (low-average versus high anxiety) as between-subjects factor, and go/nogo and exogenous/endogenous as within-subjects factors. For these analyses a significance level of *p* < .05 was used.

## 3 Results

### 3.1 Behavioral Results

The adolescents in the high anxiety group performed significantly worse on the Gambling task compared to the adolescents of the low-average anxiety group. As can be seen in Figure 2, there was a difference in the distribution of gambles/passes across trials between both anxiety groups. This distribution was quantified in a contrast measure defined as (M-S)/(S+M) where S is the proportion of gambles in the NOGO trials and M the proportion of gambles in all other trials. This contrast ranges between -1 (only gambles in the NOGO trials) and 1 (only gambles in the other trials), with 0 indicating equal percentages of gambles in both trial categories. The scatterplot in Figure 2 clearly shows that the contrast was close to 1 for the adolescents in the low-average anxiety group (*M* = 0.96, SD = 0.02), and significantly higher compared to the contrast for the high anxiety group (*M* = 0.88, SD = 0.14) (*p* < .03).

**Figure 2:**
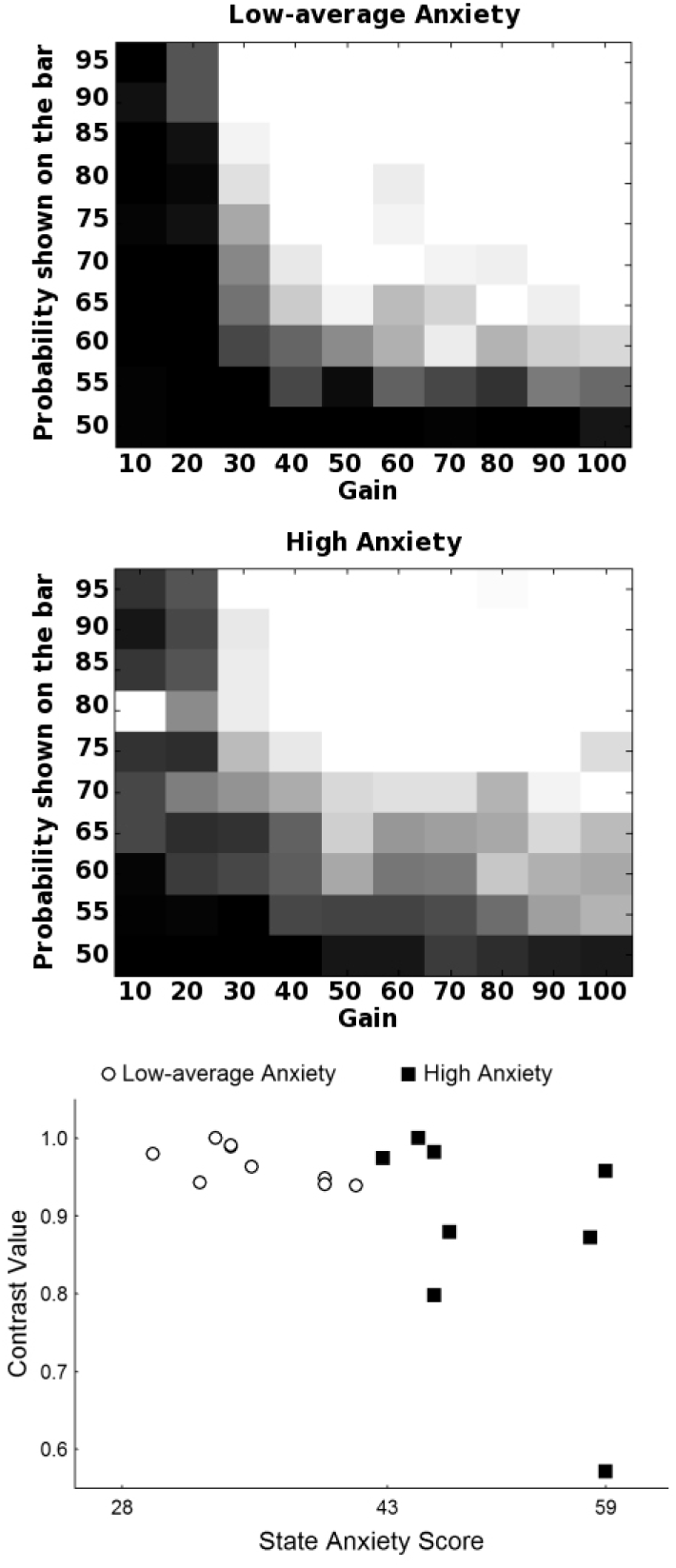
Percentage of gambles made in the different trials of the Gambling paradigm for the low-average and high anxiety group. Darker squares indicate a higher percentage of inhibitions (black = 100% pass), lighter squares indicate a higher percentage of gambles (white = 100% gamble). Trials are ordered according to the proportional division of the stimulus bar (Y-axis) and the amount of points that could be won (X-axis). The bottom graph shows the contrast values calculated based on the behavioral diagrams of each participant in relation to the maternal state anxiety score at 12–22 weeks of pregnancy

Next to a difference in this gambling contrast the adolescents in the high anxiety group also earned significantly less points across the four sessions compared to the adolescents in the low-average group (low-average: *M* = 7953, SD = 494.22; high: *M* = 6735, SD = 1460.3; *p* < .02). There was no effect of anxiety on the reaction time or the standard deviation of the reaction time.

### 3.2 fMRI Results

#### Effects of condition in the low-average anxiety group

Significant regions of activation for the effects of condition in the adolescents of the low-average anxiety group are shown in Table 1.

**Table 1:** Regions activated during the decision phase of the Gambling paradigm in the adolescents of the low-average anxiety group

|  | Side | x | y | z | *t*-value |
| --- | --- | --- | --- | --- | --- |
| **Main effect go/nogo** |  |  |  |  |  |
| go > nogo |  |  |  |  |  |
| Inferior precentral sulcus | L | -60 | 6 | 30 | 5.21 |
| Insula | L | -38 | 2 | 12 | 5.70 |
| ACC | R | 4 | -2 | 54 | 6.63 |
|  | R | 2 | -16 | 54 | 5.55 |
|  | R | 6 | 4 | 44 | 5.02 |
| Primary motor cortex | L | -48 | -20 | 56 | 9.45 |
|  | R | 42 | -20 | 56 | 10.00 |
| go < nogo |  |  |  |  |  |
| Middle frontal gyrus | L | -36 | 16 | 56 | -4.71 |
| Superior frontal gyrus | R | 22 | 24 | 58 | -4.86 |
| **Main effect of exogenous/endogenous** |  |  |  |  |  |
| Exogenous > endogenous |  |  |  |  |  |
| *a* Middle frontal gyrus | L | -30 | 32 | 48 | 6.07 |
| Exogenous < endogenous |  |  |  |  |  |
| *b* Superior frontal sulcus | R | 28 | 4 | 58 | -6.27 |
| *c* Inferior precentral sulcus | R | 54 | 12 | 26 | -4.69 |
| *d* Inferior frontal junction | R | 38 | 22 | 36 | -4.39 |
| *e* ACC | R | 6 | 20 | 48 | -9.84 |
|  | R | 10 | 30 | 34 | -7.92 |
|  | R | 0 | 18 | 54 | -6.39 |
| *f* operculum | L | -30 | 28 | 0 | -5.60 |
| *g* operculum | R | 32 | 22 | 4 | -8.69 |
| *h* Inferior frontal sulcus | R | 36 | 32 | 26 | -6.16 |

*Notes:* Activations are assessed at false discovery rate (FDR), $p < .05$ . L: Left; R: Right. Letters correspond to letters in Figure 3.

Five prefrontal clusters showed significant differences when assessing the effect of go (including the GO and GAMBLE trials) versus nogo (including the NOGO and PASS trials) decision trials. Clusters in left middle frontal gyrus and right superior frontal sulcus showed a go < nogo e ect. In contrast, ACC, left insula and left inferior precentral sulcus showed a go > nogo e ect. As expected, the strongest go > nogo effects were observed in primary motor cortex.

A number of clusters showed significant differences related to the exogenous (GO and NOGO) and endogenous (GAMBLE and PASS) trials. These clusters are shown in Figure 3. Except for one cluster in left middle frontal gyrus all clusters showed greater activations in the endogenous compared to the exogenous trials. This effect included clusters in right dorsal ACC, right inferior frontal sulcus, right inferior frontal junction, right inferior precentral sulcus, right superior frontal sulcus, and left operculum.

**Figure 3:**
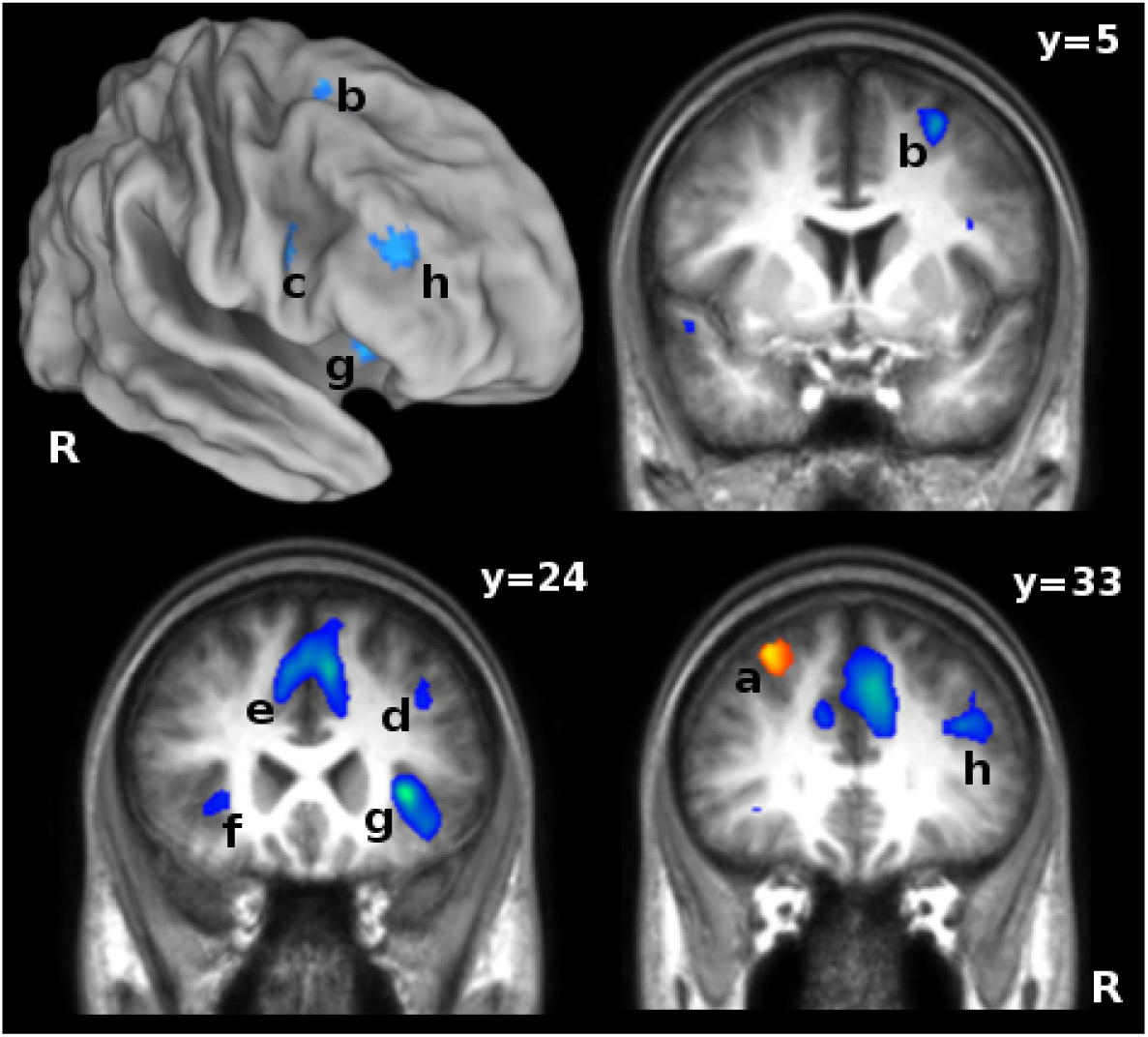
Activation maps of the exogenous/endogenous manipulation in the Gambling paradigm for the adolescents in the low-average anxiety group. Left top shows an overview of the activations plotted on the right hemisphere of the PALS-B12 atlas using CARET software (http://brainmap.wustl.edu/caret, Van Essen, 2002). Coronal slices show activations plotted on the mean T1 image of the nine adolescents of the low-average anxiety group. Maps are thresholded at FDR *p* < .05, *t* > 3.59. Red colors indicate exogenous > endogenous activations, blue indicates endogenous > exogenous activations. Letters indicate clusters as shown in Table 1.

There were no clusters that showed a go/nogo by exogenous/endogenous interaction.

#### Effects of anxiety

Comparing the BOLD response of the adolescents in the low-average anxiety group with that of the adolescents in the high anxiety group yielded 15 significant clusters in prefrontal cortex (see Table 2). Investigating the percentage of signal change in each of these ROIs, revealed that the relationship between the BOLD response and the level of antenatal maternal anxiety differed depending on the assessed ROI.

**Table 2:** Regions that are activated during the decision phase of the Gambling paradigm when comparing the adolescents of the low-average anxiety group to the adolescents of the high anxiety group

|  |  | Side | x | y | z | F-value |
| --- | --- | --- | --- | --- | --- | --- |
| **Main effect of anxiety** |  |  |  |  |  |  |
| 1 | Inferior precentral sulcus, superior | L | -37 | 8 | 50 | 8.66 |
| 2 | Inferior frontal gyrus, pars opercularis | L | -56 | 16 | 16 | 9.28 |
| 3 | Middle frontal gyrus, superior | L | -32 | 20 | 52 | 6.34 |
| 4 | ACC | R | 8 | 34 | 42 | 7.04 |
| 5 | Superior frontal junction | L | -22 | 6 | 58 | 9.50 |
| 6 | Inferior frontal sulcus | L | -36 | 10 | 32 | 10.26 |
| 7 | Inferior frontal gyrus, pars triangularis | L | -48 | 40 | 8 |  |
| **Anxiety by exogenous/endogenous interaction** |  |  |  |  |  |  |
| 8 | Superior frontal junction | L | -28 | -10 | 68 | 13.92 |
| 9 | Middle frontal gyrus, anterior | L | -24 | 56 | 26 | 8.62 |
| 10 | Inferior precentral sulcus | L | -52 | 12 | 34 | 8.85 |
| 11 | Inferior precentral sulcus | R | 58 | 10 | 30 | 10.43 |
| 12 | Inferior frontal junction | R | 44 | 20 | 40 | 6.76 |
| 13 | Middle frontal gyrus, inferior | R | 40 | 38 | 20 | 10.70 |
| **Anxiety by go/nogo interaction** |  |  |  |  |  |  |
| 14 | Superior frontal junction | R | 32 | -8 | 66 | 7.39 |
| **Anxiety by go/nogo by exogenous/endogenous interaction** |  |  |  |  |  |  |
| 15 | Superior precentral sulcus | L | -48 | 0 | 38 | 12.75 |

Notes: Activations are assessed at $p < .05$ corrected for multiple comparisons (FWE). L: Left; R: Right. Numbers correspond to numbers in Figure 4, 5 and 6.

*First*, there were six ROIs that showed a main effect of anxiety, without any interactions between anxiety and the within subject variables. These ROIs could be further divided based on the difference between both anxiety groups. Four ROIs showed activations in the adolescents of the low-average anxiety group, but not in the adolescents of the high anxiety group (see Figure 4). These included left inferior precentral sulcus (F (1,15) = 8.06; *p* < .01), left inferior frontal gyrus pars opercularis (F (1,15) = 9.5; *p* < .01), left middle frontal gyrus superior (F (1,15) = 11.5; *p* < .01) and a ROI in anterior cingulate cortex (F (1,15) = 5.8; *p* < .03). In contrast, as shown in Figure 5, left superior frontal junction (F (1,15) = 12.8; *p* < .01) and a cluster in left inferior frontal sulcus (F (1,15) = 6.99; *p* < .02) showed no activation in the low-average anxiety group, but positive activations in the high anxiety group. A similar effect was observed in a cluster in left inferior frontal gyrus pars triangularis (F (1,15) = 7.31; *p* < .02). Here the adolescents of the high anxiety group showed negative activations in absence of activations for the low-average anxiety group. This cluster also showed a significant anxiety by exogenous/endogenous interaction (F (1,15) = 4.96; *p* < .05), due to a positive activation observed in activation for the PASS trials of the low-average anxiety group.

**Figure 4:**
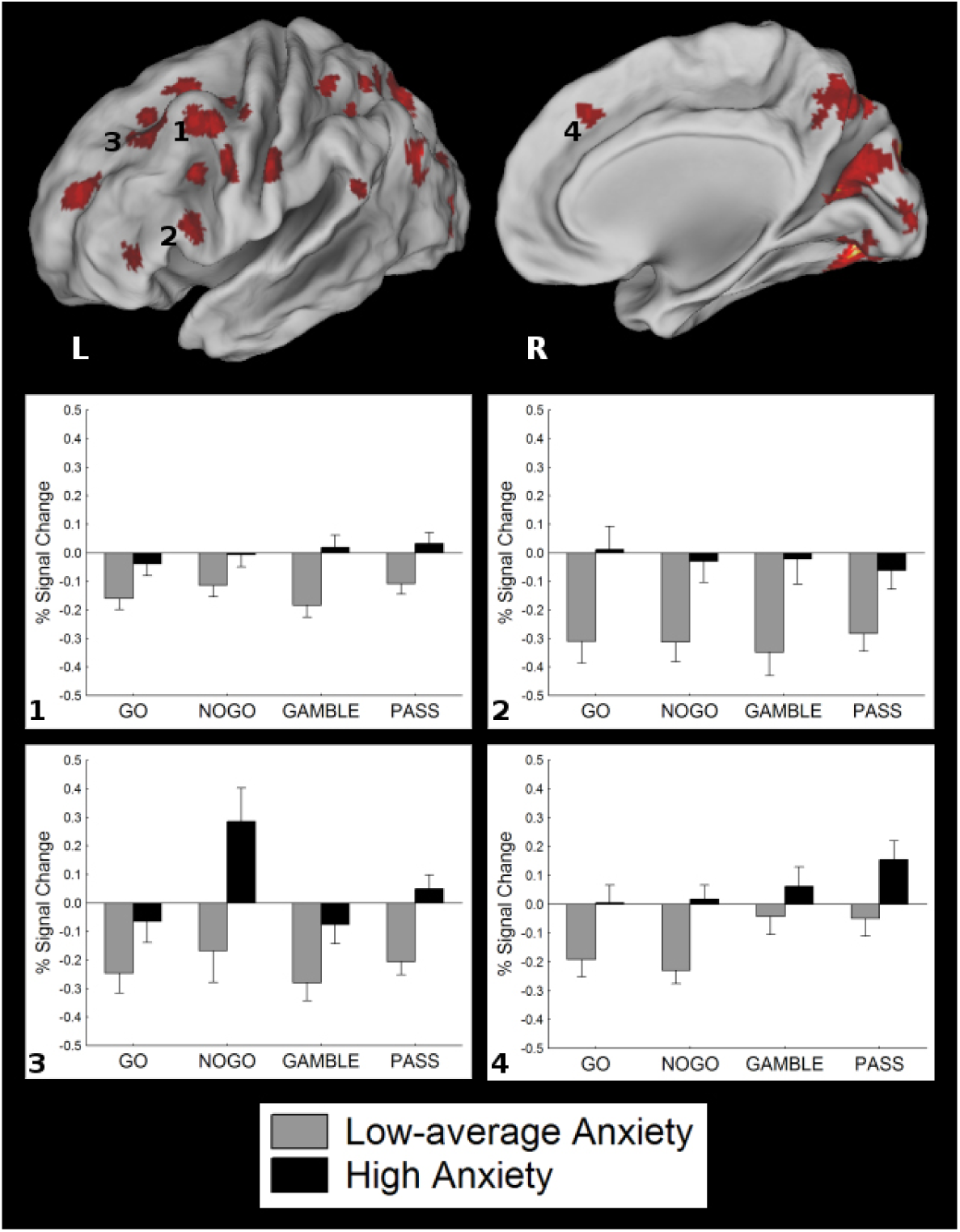
Overview of clusters showing more deactivation in the adolescents of the low-average anxiety group. Bar plots show the percentage of signal change in the designated clusters for each of the four decision conditions of the Gambling paradigm. Activated clusters are plotted on the left and right hemisphere of the PALS-B12 atlas using CARET software (http://brainmap.wustl.edu/caret, Van Essen, 2002). Activations are thresholded at *p* < .05 corrected for multiple comparisons (FWE), *F* > 4.39. Numbers indicate clusters as shown in Table 2.

**Figure 5:**
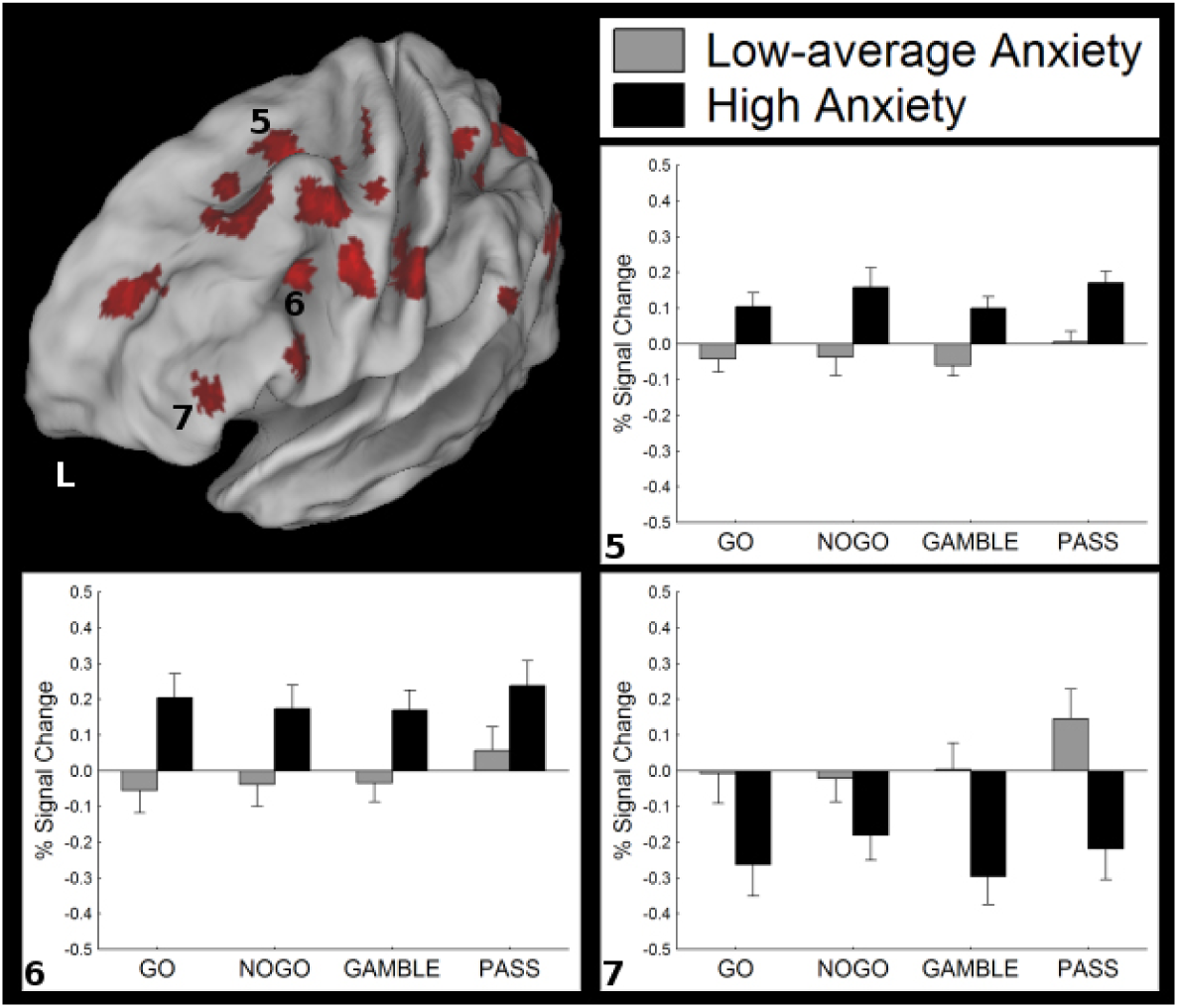
Overview of clusters showing activation in the adolescents of the high anxiety group but not in the adolescents of the low-average anxiety group. Bar plots show the percentage of signal change in the designated clusters for each of the four decision conditions of the Gambling paradigm. Activated clusters are plotted on the left hemisphere of the PALS-B12 atlas using CARET software (http://brainmap.wustl.edu/caret, Van Essen, 2002). Activations are thresholded at *p* < .05 corrected for multiple comparisons (FWE), *F* > 4.39. Numbers indicate clusters as shown in Table 2.

*Second*, Figure 6 shows six ROIs where the ANOVA yielded an anxiety by exogenous/endogenous interaction. Again these could be subdivided. In four ROIs an exogenous/endogenous modulation could be observed in the activations of the adolescents of the low-average anxiety group. However, this modulation was absent in the adolescents of the high anxiety group. These ROIs included right inferior frontal junction (F (1,15) = 11.27; *p* < .01), right middle frontal gyrus inferior (F (1,15) = 16.94; *p* < .001), and a bilateral ROI in inferior precentral sulcus (left: F (1,15) = 13.18; *p* < .01; right: F (1,15) = 6.8; *p* < .02). Next to these four clusters there were two clusters that showed an exogenous/endogenous effect in the adolescents of both anxiety groups. However, the difference in activation between the adolescents in the low-average anxiety group and those in the high anxiety group was larger in the endogenous trials compared to the difference between both groups in the exogenous trials. These clusters were found in left superior frontal junction (F (1,15) = 5.26; *p* < .04) and left middle frontal gyrus anterior (*F* (1,15) = 6.8; *p* < .02).

**Figure 6:**
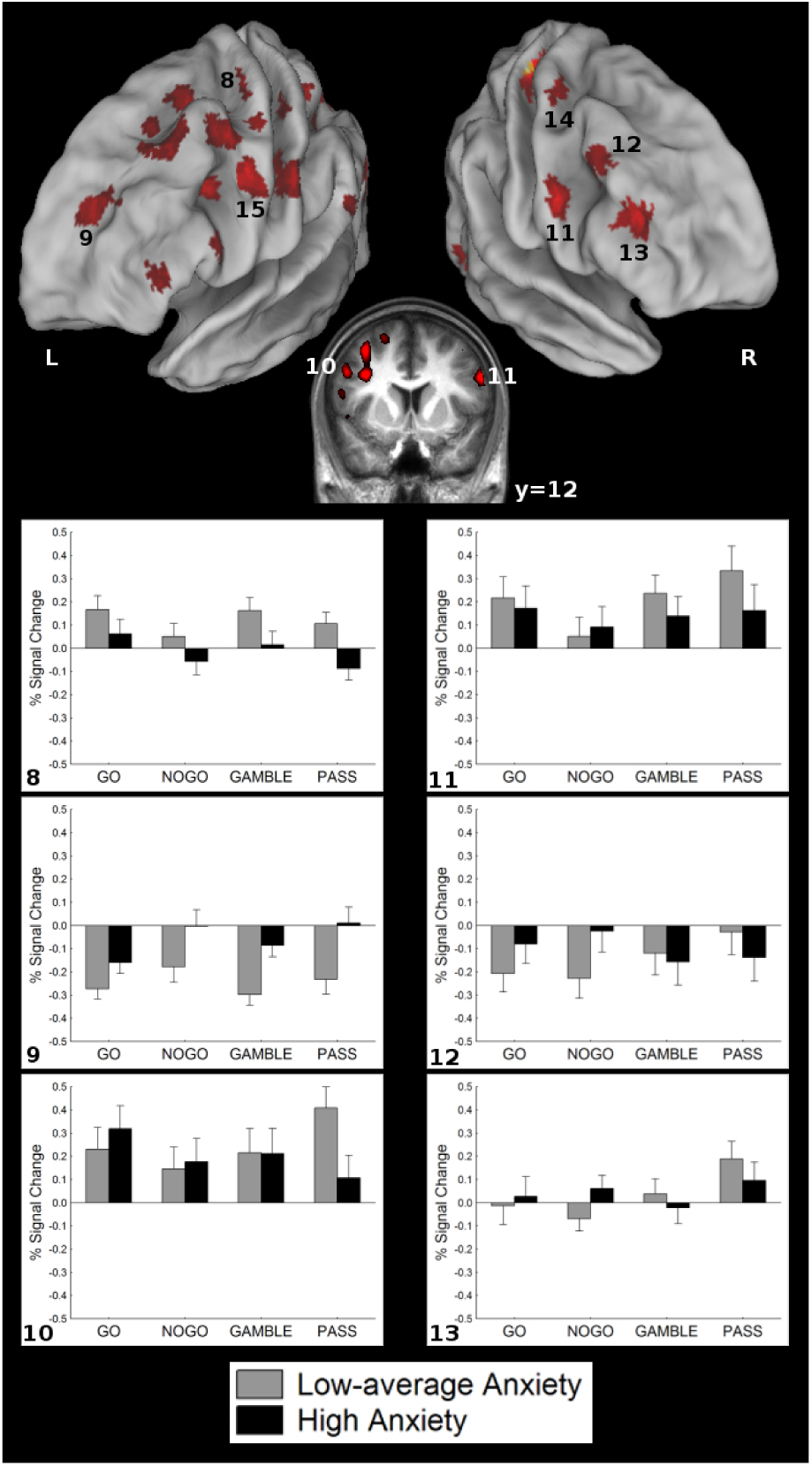
Overview of clusters showing a group by exogenous/endogenous interaction. Bar plots show the percentage of signal change in the designated clusters for each of the four decision conditions of the Gambling paradigm. Bar plots are not shown for clusters 14 and 15. Cluster 14 showed a group by go/nogo interaction, cluster 15 showed a group by go/nogo by exogenous/-endogenous interaction. Activated clusters are plotted on the left and right hemisphere of the PALS-B12 atlas using CARET software (http://brainmap.wustl.edu/caret, Van Essen, 2002). Activations are thresholded at *p* < .05 corrected for multiple comparisons (FWE), *F* > 4.39. Numbers indicate clusters as shown in Table 2.

*Third*, one cluster yielded an anxiety by go/nogo interaction. This cluster in right superior frontal junction (*F* (1,15) = 11.68; *p* < .01) was active only in the go (GO and GAMBLE) trials of the adolescents of the low-average group.

*Fourth*, there was one cluster that yielded a 3-way anxiety by go/nogo by exogenous/endogenous interaction. However, it is not clear what caused the effect in this left superior precentral sulcus cluster (*F* (1,15) = 6.42; *p* < .02), as post-hoc comparisons did not yield significant differences between both anxiety groups in each of the four conditions. Borderline differences were observed in the GO (*p* = .06) and PASS (*p* = .08) condition.

In summary, we found clusters that were active only in the adolescents of the low-average anxiety group (*n* = 4), clusters that were active only in the adolescents of the high anxiety group (*n* = 3), clusters that showed an exogenous/endogenous modulation only in the adolescents of the low-average anxiety group (*n* = 4), clusters where the difference between both anxiety groups was larger for endogenous trials (*n* = 2), one cluster that was active only in the go trials of adolescents in the low-average anxiety group and finally one cluster showing mixed results. It is important to note that most clusters were found in the left hemisphere, except for the clusters showing an exogenous/endogenous modulation in the adolescents of the low-average group but not in the adolescents of the high anxiety group.

There was no overlap between the clusters that showed anxiety related modulations and the clusters that were activated in the adolescents of the low-average anxiety group by the go/nogo condition manipulation. In contrast, there were four clusters, showing more activity in endogenous compared to exogenous trail conditions in the low-average anxiety group, that overlapped with clusters that were also significant in the comparison between the two anxiety groups. These clusters were observed in ACC, inferior frontal sulcus, inferior frontal junction and inferior precentral sulcus. All were lateralized to the right hemisphere.

## 4 Discussion

The BOLD response of 20 year old adolescent boys measured during decision making in a Gambling paradigm was found to be related to the level of anxiety reported by their mothers during weeks 12–22 of pregnancy. We observed a heterogeneous pattern of differences in brain activation related to antenatal maternal anxiety in a number of areas in prefrontal cortex. Some of these clusters were also modulated by the exogenous/endogenous cognitive control manipulation in the Gambling paradigm but only in the adolescents of the low-average anxiety group. Opposed to our previous hypothesis, that was based on purely neurocognitive behavioral results, the BOLD modulations that yielded a relationship with antenatal maternal anxiety were not confined to orbitofrontal cortex. There were no effects of antenatal maternal anxiety on brain activity related to mere response inhibition, nor in brain areas commonly involved in successful inhibition. These findings conform our previous neurocognitive behavioral results as well as our ERP results in the same sample, where no behavioral, nor brain wave differences were found in simple Go/Nogo conditions.

As hypothesized the exogenous/endogenous manipulation in the Gambling task yielded the most important modulations related to antenatal maternal anxiety. In four prefrontal clusters, including the right inferior frontal junction, an anxiety by exogenous/endogenous interaction indicated that the adolescents in the low-average anxiety group showed more positive activity in the endogenous compared to the exogenous trials, while such modulation was absent in the adolescents of the high anxiety group. Because these clusters were directly related to the endogenous aspect of the task, as revealed by the separate exogenous/endogenous contrast calculated for the adolescents of the low-average anxiety group only, it is likely that these clusters are relevant for the implementation of endogenous cognitive control. These results are in line with other studies associating inferior frontal junction with endogenous cognitive control in gambling (Zysset et al., 2006) and task-switching paradigms (Brass et al., 2005; Forstmann et al., 2005). Right inferior frontal sulcus has also been associated with set-shifting (Konishi et al., 1999) and the right hemisphere lateralization of these clusters is consistent with a PET study by Rogers et al. (1999), who found mainly right hemisphere activations when comparing decision making to a control condition. More importantly, the overlap between these clusters and clusters indicating a difference in brain activity related to antenatal maternal anxiety confirms our previous hypothesis regarding an association between antenatal maternal anxiety and endogenous cognitive control (Mennes et al., 2006; Van den Bergh et al., 2005, 2006). The observed differences in activation suggest that modulations in brain activity typically related to endogenous cognitive control are not present in the adolescents of the high anxiety group. Therefore, they suggest less efficient implementation of endogenous cognitive control in the offspring of mothers experiencing high levels of anxiety during their pregnancy.

A number of prefrontal clusters yielded a main effect of anxiety. Half of these clusters showed this effect due to significant deactivation in the adolescents of the low-average anxiety group compared to no activity at all in the high anxiety group. Deactivation of prefrontal regions including the posterior insula during cognitively demanding tasks is associated with an increase in the need for focused attention towards the specific task demands (Gusnard & Raichle, 2001; Lawrence, Ross, Hoffmann, Garavan, & Stein, 2003) or to an optimization of performance by minimizing interference from task-irrelevant areas (Tomasi, Ernst, Caparelli, & Chang, 2006). In line with this, Hester et al. (2004) have found that unsuccessful inhibition trials were associated with a failure to deactivate the insula, suggesting that a failure in attention regulation was the cause for unsuccessful performance. In the present study the insula was equally deactivated across exogenous and endogenous conditions in the adolescents of the low-average anxiety group, suggesting task related, but not condition specific deactivation. All these findings are in line with the idea that there is a ‘default network’ of brain areas that deactivates during goal-directed activities (Gusnard & Raichle, 2001). In contrast, the other clusters that yielded a main effect of anxiety showed no activation in the adolescents of the low-average anxiety group compared to either clearly positive or negative activations for the adolescents of the high anxiety group, suggesting that these adolescents effectively use other areas of prefrontal cortex during decision making compared to the adolescents in the low-average anxiety group. This is consistent with findings in children with ADHD showing that these children use a more diffuse network of prefrontal regions compared to control children (Durston et al., 2003). Similar observations were made in adults with ADHD, even after normalization of their behavior through medication (Schweitzer et al., 2004).

The behavioral results of the current follow-up phase are in line with the results obtained with the same task under EEG recordings at age 17 (Mennes et al., 2009). As can be seen in Figure 2, the adolescents in the high anxiety group showed a different pattern of decision making across the trials of the Gambling paradigm. This was quantified in the gambling contrast value that was statistically lower in the adolescents of the high anxiety group, indicating a more diverse pattern of decision making. The scatterplot in Figure 2 shows that there was considerably more variation in this contrast value for the high anxiety group compared to the values for the low-average anxiety group. In addition the adolescents in the high anxiety group earned a significantly lower amount of points. This indicates that the pattern of decision making of these adolescents is less efficient compared to the pattern observed in the adolescents of the low-average anxiety group.

Taken together these results indicate that there is a relationship between the level of anxiety experienced by a mother during pregnancy and the functional brain activity in the prefrontal cortex of her offspring. Adolescents of mothers reporting high levels of anxiety during weeks 12–22 of their pregnancy seem to show discrepant activation in areas that are recruited by adolescents of mothers reporting low to average levels of anxiety. In addition the adolescents in the high anxiety group recruited areas that were not active in the adolescents of the low-average anxiety group. The suboptimal behavioral results for this group suggest that the network recruited by adolescents in the high anxiety group is less efficient compared to the network of areas used by adolescents in the low-average anxiety group. Especially areas that are important for the implementation of endogenous cognitive control were not modulated by this type of cognitive control in the adolescents of the high anxiety group.

It is possible that a difference in maturation underlies the observed differences in activation between the adolescents in the low-average and high anxiety groups. The prefrontal cortex still undergoes developmental changes during adolescence (Paus, 2005). Imaging studies on the development of cognitive control provide evidence showing also functional changes in the involved prefrontal circuitry with maturation (Bunge, Dudukovic, Thomason, Vaidya, & Gabrieli, 2002; Durston et al., 2002). Dysmaturation of prefrontal activation has also been associated with hypofrontality that is commonly observed in children with ADHD (Booth et al., 2005; Rubia et al., 1999, 2000; Smith, Taylor, Brammer, Toone, & Rubia, 2006). However, although the majority of areas that were related to antenatal maternal anxiety showed less activation in the adolescents of the high anxiety group, we also found areas showing increased activation for this group. This suggests that the adolescents in the high anxiety group compensate for the less efficient activation of certain areas by recruiting extra areas that were not recruited by the adolescents in the low-average anxiety group. It is thus plausible that the adolescents in the high anxiety group show small maturational delays across different areas of prefrontal cortex compared to the adolescents in the low-average anxiety group. This hypothesis is in line with the hypothesis based on the ERP results with the same task (Mennes et al., 2009). In that study an increased frontal P2a peak was found in the adolescents of the high anxiety group, consistent with higher P2a peak values in younger participants (Jonkman, 2006).

The current results are the first to show a relationship between functional brain activity as measured with fMRI and the level of anxiety experienced by a mother during pregnancy. A recent study by Buss et al. reported on structural effects as they found that an adverse prenatal environment negatively influenced hippocampal volume in the absence of a positive postnatal environment (Buss et al., 2007). This effect was only observed in girls. Structural effects of the prenatal environment have been extensively documented in animal research (for a review see Weinstock, 2001). These studies suggest that subtle developmental alterations in structures connected to the prefrontal cortex might underlay the functional differences related to antenatal maternal anxiety observed in the current study. Indeed, the current results were related to weeks 12–22 of pregnancy which is an important period for the structural differentiation of several structures that share intensive connections with to the prefrontal cortex (e.g., hippocampus, amygdala, anterior cingulate cortex, brainstem, and basal ganglia) (Garel, 2004; Levitt, 2003; Nowakowski & Hayes, 2002). However, there is no strict relationship between abnormal brain structure and dysfunctional behavior. It can even be the case that dysfunctional behavior is the cause of structural modifications (Rubia, 2002).

Opposed to our previous hypothesis that the functioning of the orbitofrontal cortex is most affected by antenatal maternal anxiety, we did not observe any anxiety related activation differences in that part of prefrontal cortex. This hypothesis was based on a review of imaging studies that used tasks that were similar to the ones we used. However, the Gambling paradigm that was used here, although argued to tap into the same underlying functions as the task-switching paradigm on which we based our hypothesis, was not incorporated in that review (Mennes et al., 2006). It is likely that such gambling paradigms activate different specific areas of prefrontal cortex compared to the studies included in our review. Accordingly, the current study yielded no activations related to decision making in the orbitofrontal cortex. In contrast, Rogers et al. (2004) found orbitofrontal cortex to be activated when deliberating over a decision involving large gains compared to a decision involving small gains (Rogers et al., 2004). One could also argue that the absence of effects in orbitofrontal cortex is due to the fact that the imaging sequence we used was not optimized for imaging orbitofrontal cortex. Such special sequences are needed since this part of the brain is vulnerable for imaging artifacts due to its proximity to the nasal cavity inducing artifacts by magnetization inhomogeneities between air and brain tissue. Recently techniques for optimizing signal obtained from orbitofrontal cortex have been proposed (Deichmann, Gottfried, Hutton, & Turner, 2003; Weiskopf, Hutton, Josephs, Turner, & Deichmann, 2007). However, these methods are based on a different angle of image acquisition, causing other areas of the brain to fall outside the acquisition window. Although we cannot exclude the influence of such artifacts, they are an unlikely cause for the absence of effects in orbitofrontal cortex in the current study, since these artifacts mostly occur in ventromedial orbitofrontal cortex and at the frontal pole. In contrast the areas involved in our hypothesis were situated in dorsolateral orbitofrontal cortex.

An important limitation for our conclusions is the rather limited number of participants we were able to include. Therefore we chose a fixed-effects analysis for comparing the brain activity of the adolescents of both anxiety groups. Although providing valid results for the investigated sample this type of analysis hampers possible interpretations with respect to the total population. The smaller sample size also increases the chance for type II errors, indicating that significant differences were missed. This could for instance be the case for activations in orbitofrontal cortex. However, it should be noted that due to the longitudinal design of the study it was impossible to include new participants as we would have no access to data on the level of anxiety experienced by the mothers gathered during their pregnancy. The longitudinal design is on the other hand, also an obvious strength of the current study. Within the same sample we found evidence for an impairment in endogenous cognitive control related to the level of exposure to antenatal maternal anxiety at different moments during development and using different techniques (Mennes et al., 2009; Van den Bergh et al., 2005, 2006; Mennes et al., 2006). It is unlikely that these corroborating effects are just found by chance in each of those studies.

In conclusion, the current results provide evidence for an association between the level of anxiety experienced by a mother during weeks 12–22 of her pregnancy and the functional activation of prefrontal cortex. The modulation of activity in areas that are important for the implementation of endogenous cognitive control provides compelling support for our previous hypothesis that high levels of maternal anxiety during pregnancy are related to a deficit in endogenous cognitive control. Further research into the mechanisms underlying this relationship is needed. The current results suggest that the maturation of prefrontal cortex should receive special interest.

## Acknowledgments

The authors wish to thank the participating adolescents and their families for their continuing interest in the study. We thank Ron Peeters, Silvia Kovacs and Caroline Sage for technical support during scanning. This work was supported by the Research Foundation Flanders (FWO) (#G.0211.03) and by the K.U.Leuven (IMPH/06/GHW and IDO 05/010 EEG-fMRI). LL is holder of the ‘UCB Chair on Cognitive Dysfunctions in Childhood’ at the K.U.Leuven.

## References

Booth, J. R., Burman, D. D., Meyer, J. R., Lei, Z., Trommer, B. L., Davenport, N. D., … Mesulam, M. (2005). Larger deficits in brain networks for response inhibition than for visual selective attention in attention deficit hyperactivity disorder (adhd). The Journal of Child Psychology and Psychiatry, 46, 94–111.

Brass, M., Derrfuss, J., Forstmann, B., & von Cramon, D. (2005). The role of the inferior frontal junction area in cognitive control. Trends in Cognitive Sciences, 9, 314–316.

Brass, M., & von Cramon, Y. (2004). Decomposing components of task preparation with functional magnetic resonance imaging. Journal of Cognitive Neuroscience, 16, 609–620.

Bunge, S., Dudukovic, N., Thomason, M., Vaidya, C., & Gabrieli, J. (2002). Immature frontal lobe contributions to cognitive control in children: Evidence from fmri. Neuron, 33, 301–311.

Buss, C., Lord, C., Wadiwalla, M., Hellhammer, D. H., Lupien, S. J., Meaney, M. J., & Pruessner, J. C. (2007). Maternal Care Modulates the Relationship between Prenatal Risk and Hippocampal Volume in Women But Not in Men. The Journal of Neuroscience, 27, 2592–2595.

Deichmann, R., Gottfried, J., Hutton, C., & Turner, R. (2003). Optimized EPI for fMRI studies of the orbitofrontal cortex. NeuroImage, 19, 430–441.

Durston, S., Thomas, K. M., Yang, Y., Ulug, A. M., Zimmerman, R. D., & Casey, B. (2002). A neural basis for the development of inhibitory control. Developmental Science, 5, F9–F16.

Durston, S., Tottenham, N., Thomas, K., Davidson, M., Eigsti, I., Yang, Y., … Casey, B. (2003). Differential patterns of striatal activtion in young children with and without ADHD. Biological Psychiatry, 53, 871–878.

Forstmann, D., Brass, M., Koch, I., & von Cramon, D. (2005). Internally generated and directly cued task sets: an investigation with fMRI. Neuropsychologia, 6, 943–952.

Garel, C. (2004). Mri of the fetal brain. normal development and cerebral pathologies. Berlin-Heidelberg (Germany): Springer-Verlag.

Gusnard, D., & Raichle, M. (2001). Searching for a baseline: Functional imaging and the resting human brain. Nature Reviews Neuroscience, 2, 685–694.

Hester, R. L., Murphy, K., Foxe, J. J., Foxe, D. M., Javitt, D. C., & Garavan, H. (2004). Predicting Success: Patterns of Cortical Activation and Deactivation prior to Response Inhibition. Journal of Cognitive Neuroscience, 16, 776–785.

Jonkman, L. (2006). The development of preparation, conflict monitoring and inhibition from early childhood to young adulthood; a Go/Nogo ERP study. Brain Research, 1097, 181–193.

Konishi, S., Nakajima, K., Uchida, I., Kikyo, H., Kameyama, M., & Miyashita, Y. (1999). Common inhibitory mechanism in human inferior prefrontal cortex revealed by event-related functional MRI. Brain, 122(5), 981–991.

Lawrence, N. S., Ross, T. J., Hoffmann, R., Garavan, H., & Stein, E. A. (2003). Multiple Neuronal Networks Mediate Sustained Attention. Journal of Cognitive Neuroscience, 15, 1028–1038.

Levitt, P. (2003). Structural and functional maturation of the developing primate brain. Journal of Pediatrics, 143, 35–45.

Mennes, M., Stiers, P., Lagae, L., & Van den Bergh, B. (2006). Long-term cognitive sequelae of antenatal maternal anxiety: involvement of the orbitofrontal cortex. Neuroscience & Biobehavioral Reviews, 30, 1078–1086.

Mennes, M., Van den Bergh, B., Lagae, L., & Stiers, P. (2009). Developmental brain alterations in 17 year old boys are related to antenatal maternal anxiety. Clinical Neurophysiology, 120(6), 1116–1122.

Mennes, M., Wouters, H., Van den Bergh, B., Lagae, L., & Stiers, P. (2008). Detection and resolution of conflict: Erp correlates of complex human decision making. Psychophysiology, 45(5), 714–720.

Miller, E., & Cohen, J. (2001). An integrative theory of prefrontal cortex function. Annual Review of Neuroscience, 24, 167–202.

Nowakowski, R., & Hayes, N. (2002). General principles of CNS development. In M. Johnson, Y. Munakata, & R. Gilmore (Eds.), Brain development and cognition. A reader (2nd ed., pp. 57–82). Malden, MA: Blackwell Publishers.

Paus, T. (2005). Mapping brain maturation and cognitive development during adolescence. Trends in Cognitive Sciences, 9, 60–68.

Rogers, R., Ramnani, N., Mackay, C., Wilson, J., Jezzard, P., Carter, C., & Smith, S. (2004). Distinct portions of anterior cingulated cortex and medial prefrontal cortex are activated by reward processing in separable phases of decision-making cognition. Biological Psychiatry, 55, 594–602.

Rubia, K. (2002). The dynamic approach to neurodevelopmental psychiatric disorders: use of fMRI combined with neuropsychology to elucidate the dynamics of psychiatric disorders, exemplified in ADHD and schizophrenia. Behavioural Brain Research, 130, 47–56.

Rubia, K., Overmeyer, S., Taylor, E., Brammer, M., Williams, S., A., S., … Bullmore, E. (1999). Hypofrontality in attention deficit hyperactivity disorder during higher order motor control: a study with functional MRI. The American Journal of Psychiatry, 156, 891–896.

Rubia, K., Overmeyer, S., Taylor, E., Brammer, M., Williams, S., Simmons, A., … Bullmore, E. (2000). Functional frontalisation with age: mapping neurodevelopmental trajectories with fMRI. Neuroscience & Biobehavioral Reviews, 24, 13–19.

Schweitzer, J., Lee, D., Hanford, R., Zink, C., Ely, T., Tagamets, M., … Kilts, C. (2004). Effect of methylphenidate on executive functioning in adults with attention-deficit/hyperactivity disorder: Normalization of behavior but not related brain activity. Biological Psychiatry, 56, 597–606.

Smith, A. B., Taylor, E., Brammer, M., Toone, B., & Rubia, K. (2006). Task-Specific Hypoactivation in Prefrontal and Temporoparietal Brain Regions During Motor Inhibition and Task Switching in Medication-Naive Children and Adolescents With Attention Deficit Hyperactivity Disorder. American Journal of Psychiatry, 163, 1044–1051.

Tomasi, D., Ernst, T., Caparelli, E., & Chang, L. (2006). Common deactivation patterns during working memory and visual attention tasks: An intra-subject fMRI study at 4 Tesla. Human Brain Mapping, 27, 694–705.

Van den Bergh, B., Mennes, M., Oosterlaan, J., Stevens, V., Stiers, P., Marcoen, A., & Lagae, L. (2005). High antenatal maternal anxiety is related to impulsivity during performance on cognitive tasks in 14- and 15-year-olds. Neuroscience & Biobehavioral Reviews, 29, 259–269.

Van den Bergh, B., Mennes, M., Stevens, V., Van der Meere, J., Börger, N., Stiers, P., … Lagae, L. (2006). ADHD deficit as measured in adolescent boys with a continuous performance task is related to antenatal maternal anxiety. Pediatric Research, 59, 78–82.

Van den Bergh, B., Van Calster, B., Smits, T., Van Huffel, S., & Lagae, L. (2008). Antenatal Maternal Anxiety is Related to HPA-Axis Dysregulation and Self-Reported Depressive Symptoms in Adolescence: A Prospective Study on the Fetal Origins of Depressed Mood. Neuropsychopharmacology, 33, 536–545.

Van Essen, D. (2002). Windows on the brain. The emerging role of atlases and databases in neuroscience. Current Opinion in Neurobiology, 12, 574–579.

Weinstock, M. (2001). Alterations induced by gestational stress in brain morphology and behaviour of the offspring. Progress in Neurobiology, 65, 427–451.

Weiskopf, N., Hutton, C., Josephs, O., Turner, R., & Deichmann, R. (2007). Optimized EPI for fMRI studies of the orbitofrontal cortex: compensation of susceptibility-induced gradients in the readout direction. Magnetic Resonance Materials in Physics, Biology and Medicine, 20, 39–49.

Zysset, S., Wendt, C., Volz, K., Neumann, J., Huber, O., & von Cramon, D. (2006). The neural implementation of multi-attribute decision making: A parametric fMRI study with human subjects. NeuroImage, 31, 1380–1388.

